# ReSCU-Nets: recurrent U-Nets for segmentation of multidimensional microscopy data

**DOI:** 10.1101/2024.11.28.625889

**Authors:** Raymond Hawkins, Negar Balaghi, Katheryn E. Rothenberg, Michelle Ly, Rodrigo Fernandez-Gonzalez

## Abstract

Segmenting multi-dimensional microscopy data requires high accuracy across many images (e.g. timepoints or Z slices) and is thus a labour-intensive part of biological image processing pipelines. We present ReSCU-Nets, recurrent convolutional neural networks that use the segmentation results from the previous frame as a prompt to segment the current frame. We demonstrate that ReSCU-Nets outperform state-of-the-art image segmentation models in different segmentation tasks on time-lapse microscopy sequences.

## MAIN

Quantification of microscopy images facilitates detection and reproducibility of subtle phenotypes in cell and developmental biology^1^. Image quantification pipelines often begin with segmentation of the structures to be measured, such as organelles, cells, or cell ensembles. If conducted manually, segmentation is labor-intensive and prone to artefacts and human bias, especially for multidimensional images (e.g. time-lapse data) in which many frames must be segmented. Neural network architectures, such as the U-Net^2,3^, have improved the accuracy of automated image segmentation. However, segmentation of multidimensional images, particularly timeseries, remains challenging. Photobleaching and environmental changes can reduce the signal-to-noise ratio during image acquisition. Furthermore, biological structures can change their morphology or molecular composition over time, limiting the ability of neural networks to recognize the same object in multiple images of a sequence^4^.

The sequential information in timelapse images provides an avenue for improving segmentation. A common way to use temporal information is to add recurrence^5^. In a recurrent network, the output of a network layer depends on both the current input and previous outputs. The Long Short-Term Memory U-Net (LSTM U-Net) is a U-Net in which conventional convolutional layers are replaced with convolutional LSTM layers that have the ability to recall information previously seen during inference^5–7^. LSTM U-Nets provide improved accuracy over U-Nets for timelapse microscopy segmentation^5^. However, because recurrence in LSTM U-Nets is within network layers, the segmentation results are solely based on the image being considered, reducing performance when signal-to-noise is low. Promptable segmentation methods, such as the Segment Anything Model (SAM)^8^, can be used recurrently to provide the network with additional information. In this approach, the network inputs are the image and one or several prompts that describe the locations of objects in the form of coordinates, bounding boxes, or masks. The user can provide the network with prompts for the first image of a sequence, and the segmentation masks produced by the network can be used as prompts for subsequent images in the sequence^9^. SAM is built on large transformer models that require an abundance of training data (over 1 billion masks)^8^, something not accessible to the standard microscopist.

We set out to design a neural network architecture to accurately segment multidimensional microscopy images with limited training data. Thus, we began with a U-Net architecture, given the low data requirements of U-Net training^2,3^. To incorporate temporal context in the U-Net, we added a prompt encoder (Figure 1A-B). The prompt encoder used the output mask produced by the network to inform the segmentation of the same object in the following timeframe. The image to be segmented and the prompt were fed into the network through split input streams that were eventually concatenated. We refer to this architecture as the Recurrent Split Concatenated U-Net (ReSCU-Net).

**Figure 1.**
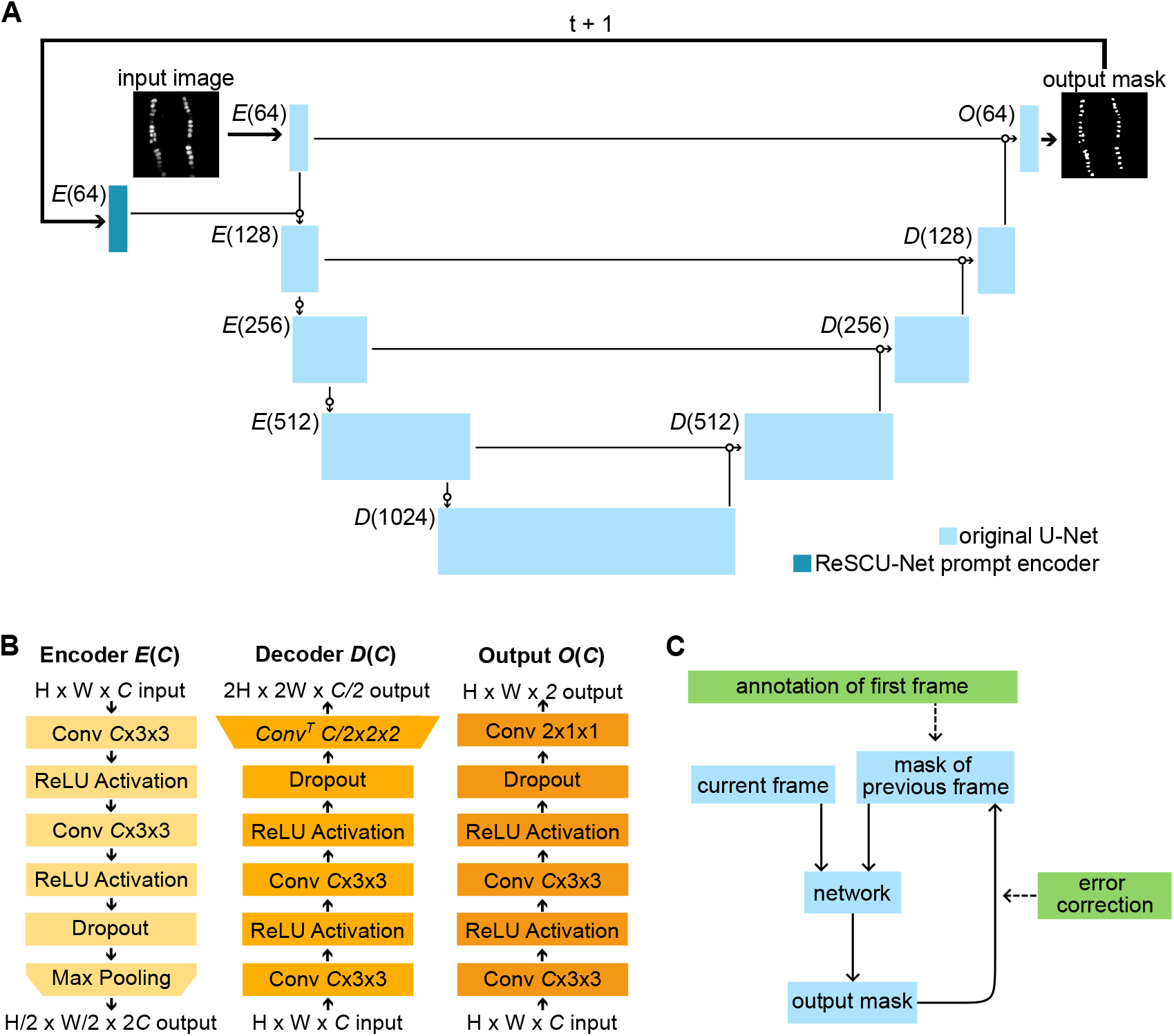
ReSCU-Nets: recurrent neural networks for multidimensional image segmentation and tracking. **(A)** ReSCU-Net architecture showing the original U-Net backbone (light blue) and the ReSCU-Net prompt encoder (dark blue). **(B)** Layer diagram of the encoder (light yellow), decoder (dark yellow), and output blocks (orange). **(C)** Example workflow for segmenting movies with ReSCU-Nets showing automated components (blue) and user interventions (green).

To facilitate ReSCU-Net design and assess network performance, we assembled three timelapse datasets, each representing unique segmentation and tracking challenges^10^. Datasets were generated by imaging living *Drosophila* embryos, and included fluorescent reporters for the nuclei of a subset of cardiac progenitors (cardioblasts) as they migrate to form the heart tube^11^ and the adherens junctions of epidermal cells migrating to repair wounds^12^. In the cardioblast dataset, the goal was to segment and track each nucleus to reconstruct the dynamics of heart development (Supplementary Figure 1A and Supplementary Video 1). Two epidermal datasets were used, one to investigate wound closure dynamics by segmenting the wound margin (Supplementary Figure 1B and Supplementary Video 2), and a second to study the cell movements and shape changes associated with the wound response by delineating the cells around the wound (Supplementary Figure 1C and Supplementary Video 3).

To determine the optimal prompt encoder architecture, we trained networks on each dataset using prompt encoder depths varying from 0 (no prompt encoder, the previous mask and the current image are concatenated before entering the network) to 4 (there are as many prompt encoder blocks as there are image encoder blocks) (Supplementary Figure 2A). To evaluate network performance, we matched each object (nucleus, wound margin, or cell) in the ground truth of an image with the object in the network output for that image with the highest intersection over union (IOU). For cardioblast nuclei and epidermal cells, the choice of prompt encoder depth affected the segmentation IOU by less than 1% (Supplementary Figure 2B, D). In contrast, when trying to segment the wound margin, using prompt encoder depths of 1, 2 or 3 produced IOUs that were at least 10% better than those obtained with depths of 0 or 4 (*P* < 0.001, Supplementary Figure 2C). These results suggest that learning the relationships between consecutive images too early or too late are both detrimental for optimal training. Despite performing similarly, the number of trainable parameters for prompt encoder depths of 2 and 3 increased by 1.4% and 7.1%, respectively, compared to a depth of 1 (Supplementary Figure 2E), with the consequent increase in training cost. Thus, we chose a prompt encoder with one encoder block for all ReSCU-Nets, as this architecture achieved high performance with minimal network size (Figure 1A-B).

ReSCU-Nets combine the advantages of recurrent neural networks and promptable segmentation methods. To use a ReSCU-Net, the network is prompted with a segmentation of the first image in the sequence, produced manually, with a U-Net, or with other methods (Figure 1C). The network then uses its own outputs as prompts to segment the rest of the images in the sequence, with the option for human-in-the-loop error correction (Figure 1C). To facilitate use, we implemented ReSCU-Nets in PyJAMAS, an open-source image analysis tool that we develop^13^. PyJAMAS provides a user interface for training networks, and both automated and semi-automated tools to generate the initial ReSCU-Net prompts.

We compared the performance of ReSCU-Nets to U-Nets, LSTM U-Nets, and the pretrained SAM *ViT Huge*^8^. To measure the accuracy of network segmentations we calculated the IOU between the ground truth and corresponding network outputs. To assess the amount of human intervention required to annotate an image sequence, we quantified the percentage of network predictions that were true positives (IOU ≥ 75%, required no intervention), false negatives (IOU < 75%, required correction by user), and false positives (not present in the ground truth, required deletion by user). For SAM and ReSCU-Nets, false negatives were replaced with the ground truth when prompting the network for the next image in the sequence, simulating a workflow with interactive error correction (Figure 1C).

U-Nets, which are not prompted and do not consider temporal information, had the worst performance overall. Despite achieving a high accuracy on cardioblast nuclei segmentation, with an IOU of 87 ± 13% (mean ± standard deviation) (Figure 2A, D), U-Nets also segmented nuclei that were not included in the ground truth due to their low level of expression of the fluorescent reporter, indicative of a different cell fate (Figure 2A, cyan). Thus, U-Nets displayed a relatively high false positive rate at 11 ± 10%, and low true positive value of 85 ± 10% (Figure 2H). U-Nets also produced the least accurate segmentations when applied to the wound margin and epidermal cell datasets, which introduce the challenges of high photobleaching and high object density, respectively (Figure 2B). The true positive value of U-Nets was 67 ± 29% for wound margins, and 14 ± 3% for epidermal cells (Figure 2E-F, H-I). Incorporating temporal information in the form of LSTM U-Nets did not significantly improve the fidelity of segmentations for cardioblast nuclei or wound margins (Figure 2A-B, G-H). LSTM U-Nets did improve the true positive rate for epidermal cells to 37 ± 11% (*P* < 0.05, Figure 2I), but the segmentation accuracy of 49 ± 37% was inconsistent and did not represent an improvement over U-Nets (*P* > 0.19, Figure 2F). Overall, LSTM U-Nets did not significantly improve IOUs with respect to other networks (Figure 2D-F) or prevent false positives (Figure 2G, I), thus suggesting that adding temporal context in the form of LSTM layers was not sufficient to improve U-Net performance on time-lapse microscopy data.

**Figure 2.**
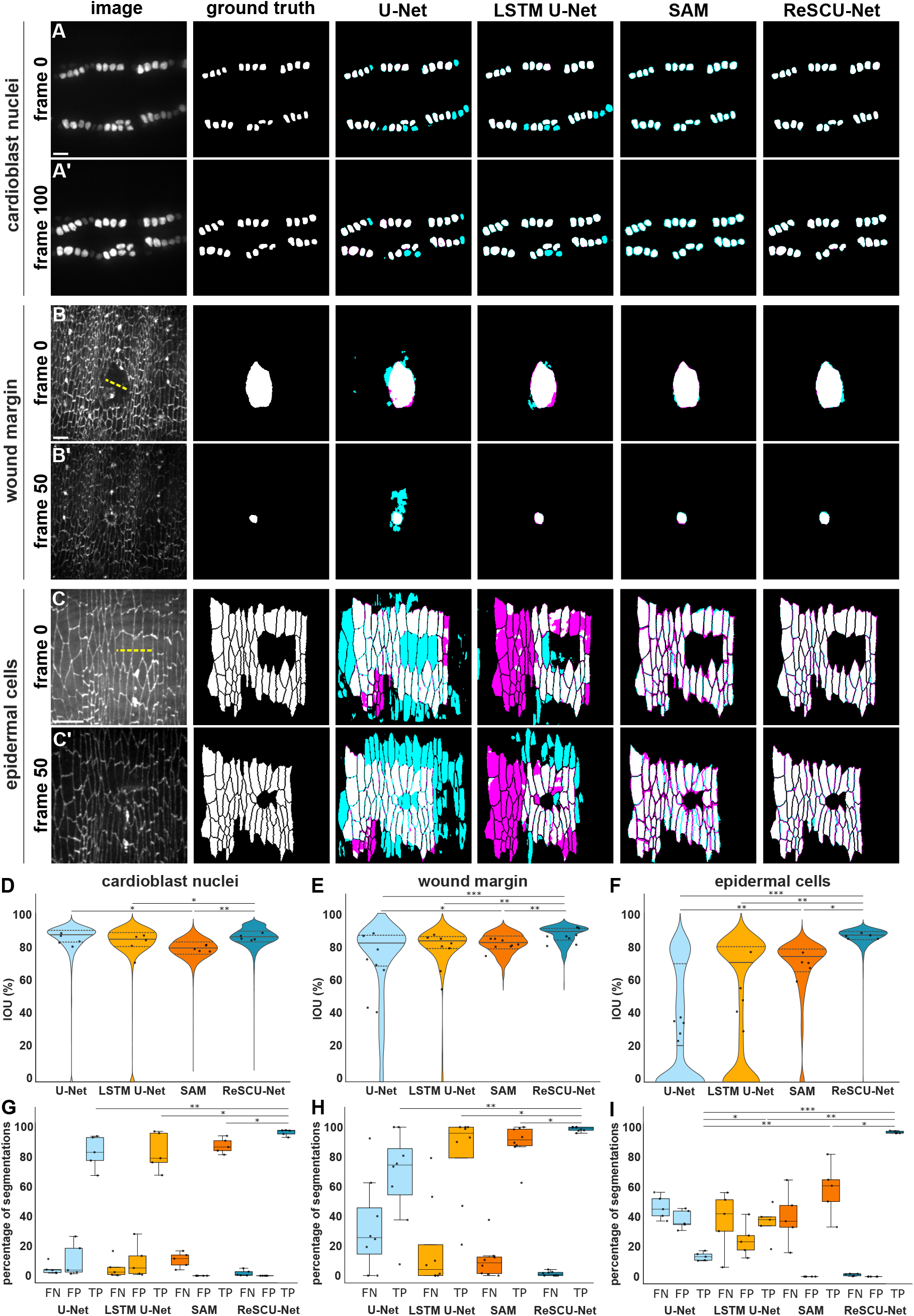
ReSCU-Nets outperform state-of-the-art networks for segmenting image sequences. **(A-C)** Images (leftmost column), ground truth segmentations, and corresponding segmentations produced by U-Net, LSTM U-Net, SAM, and ReSCU-Net (rightmost column) for time-lapse datasets depicting the nuclei of migrating cardioblasts expressing Mid^E19^:GFP (A), closure of a wound in an embryo expressing E-cadherin:tdTomato (B), and closure of a wound in an embryo expressing E-cadherin:GFP (C). Yellow dashed lines indicate the region targeted for wounding. Cyan and magenta represent pixels erroneously labelled as foreground and background, respectively. Frame number in the corresponding sequence is indicated. Anterior, left. Bars, 10 μm. **(D-F)** Test IOUs for segmentations of cardioblast nuclei (D), wound margins (E), and epidermal cells (F). Violins indicate distributions of individual object results, solid lines show the median, dashed lines indicate the quartiles, dots show movie means. **(G-I)** Percentage of segmented objects that are false negatives (FN), false positives (FP), and true positives (TP) for cardioblast nuclei (G), wound margins (H), and epidermal cells (I). Boxes indicate the quartiles, lines show the median, bars indicate 1.5x the interquartile range. (D-I) U-Net (light blue), LSTM U-Net (yellow), SAM (orange), and ReSCU-Net (dark blue). Dots show movie means. * *P* < 0.05, ** *P* < 0.01, *** *P* < 0.001. For box plots, only statistical comparisons between true positive values are shown.

We used SAM to investigate how the segmentation results change when adding temporal context in the form of prompts from previous images in the sequence (Figure 2A-C). Similar to LSTM U-Nets, SAM improved slightly (but not significantly) the true positive values of nuclear and wound margin segmentations to 89 ± 5% and 90 ± 11%, respectively (Figure 2G, H). SAM did significantly improve the results on the epidermal cell dataset with respect to U-Nets, with an IOU of 70 ± 19% (*P* < 0.01, Figure 2F) and true positive value of 60 ± 16% (*P* < 0.01, Figure 2I); a greater effect than LSTM U-Nets. Prompting also led to the disappearance of false positives in all tasks (Figure 2G, I). However, for both nuclear and wound margin datasets, SAM did not improve the accuracy of the segmentations, which in fact decreased with respect to U-Nets (*P* < 0.05, Figure 2D-E). The reduction in accuracy when using SAM was likely due to the use of a pre-trained model. SAM requires very large amounts of training data, and it is therefore unfeasible for many applications to train individual SAMs for specific tasks.

ReSCU-Nets combine the advantages of low data requirements for training and temporal context in the form of recurrent prompting. We found that ReSCU-Nets created the most accurate segmentations for all three datasets, with IOUs of 88 ± 6% (cardioblast nuclei), 90 ± 5% (wound margin), and 89 ± 6% (epidermal cells) (Figure 2A-F). Like SAM, ReSCU-Nets did not produce false positives due to prompting. The high accuracy of segmentations combined with the lack of false positives resulted in true positive values of 98 ± 2%, 99 ± 1%, and 99 ± 1% for nuclei, wound margins, and cells, respectively; significantly higher than all other networks (*P* < 0.05, Figure 2G-I). Thus, ReSCU-Nets outperform state-of-the-art models for segmentation of time-lapse microscopy data.

The success of ReSCU-Nets demonstrates the power of human-in-the-loop neural networks. Fully automated tools such as U-Nets and LSTM U-Nets often require significant manual corrections, particularly when applied to multidimensional image data. Promptable methods partially solve this problem by focusing segmentation efforts on important objects^8,9^. However, the most successful promptable methods are data-hungry^8^, and therefore are not feasible for applications in which there is limited data available for training. ReSCU-Nets improve upon existing methods by combining the low data requirements of U-Nets with recurrent prompting. Thus, despite the perceived convenience of fully automated pipelines, we demonstrate that a well-conceived semi-automated tool can maximize accuracy and minimize user intervention.

## ACKNOWLEDGMENTS

We are grateful to Veronica Castle and Willow Peterson for comments on the manuscript, and to Sergey Plotnikov and José Zariffa for useful discussions. RH was supported by a graduate scholarship from the Natural Sciences and Engineering Research Council of Canada and an Ontario Graduate Scholarship. NB was supported by an Ontario Graduate Scholarship. KR was supported by postdoctoral fellowships from the Canadian Institutes of Health Research and the Ted Rogers Centre for Heart Research. This work was funded by grants to RFG from the Natural Sciences and Engineering Research Council of Canada (418438-13), the Canada Foundation for Innovation (30279), the Translational Biology and Engineering Program of the Ted Rogers Centre for Heart Research, and the Canadian Institutes of Health Research (156279 and 186188). RFG is the Canada Research Chair in Quantitative Cell Biology and Morphogenesis.

## METHODS

### Fly stocks

*Drosophila* lines were maintained at 18°C, 25°C, or 29°C, using fly food produced in a central kitchen operated by H. Lipshitz. Before experiments, adult flies were moved to plastic cages at 25°C and embryos were collected overnight on apple juice-agar plates. Stage 14-15 embryos (12-14 hours after egg laying) of both sexes were used for all datasets. The lines used for imaging were *mid-mid*^E19^:GFP^14^, *endo-DE-cadherin:*tdTomato^15^, and *endo-DE-cadherin:*GFP^15^.

### Image acquisition

Embryos were dechorionated for 2 minutes in 50% bleach, rinsed, and staged under a dissection microscope on apple juice-agar pads. We attached embryos to a coverslip using heptane glue, dorsal side down (for imaging cardiac progenitors) or ventral side down (for imaging epidermal cells). Glued embryos were covered with a 1:1 mixture of halocarbon oil 27 and 700 (Sigma-Aldrich, St. Louis, MO)^17^. To wound the ventral epithelium for the wound margin and epidermal cell datasets, we irradiated cells with a pulsed Micropoint N2 laser (Andor, Belfast, UK) tuned to 365 nm, delivering 120 μJ pulses, 2-6 ns each. We delivered 10 consecutive pulses at discrete points 2 μm apart along a 12-14 μm line, targeting 4-5 cell areas. We imaged embryos with a Revolution XD spinning disk confocal microscope equipped with an iXon Ultra 897 camera (Andor), using a 60x oil immersion lens (Olympus, Shinjuku, Japan; NA 1.35). We acquired 16-bit Z-stacks in 0.75 μm steps with 21-27 slices per stack every 15 s (for cardioblasts), or in 0.50 μm steps with 11 slices per stack every 30 s (for epidermal cells). We used maximum intensity projections of the Z-stacks at each time point for network training and testing.

### Dataset splits

We sorted movies into train and test sets, such that the test set was at least 25% of the size of the train set, with a minimum of 5 movies in the test set (Supplementary Figure S1D-F). The cardioblast nuclei dataset contained 15 movies representing 367 unique nuclei, for a total of 71,904 individual ground truth masks. The wound margin dataset contained 39 movies, with one wound per movie, for a total of 4,319 individual masks. Lastly, the epidermal cell outline dataset contained 15 movies representing 675 unique cells, for a total of 32,120 individual masks.

### Ground truth segmentation and tracking

Ground truth segmentations were generated using a variety of manual and semi-automated methods implemented in the image analysis tools PyJAMAS^13^ and SIESTA^16^. Interactive annotations were conducted using the LiveWire method, which uses Dijkstra’s minimal cost path algorithm^18^ to identify the brightest pixel path connecting two manually selected pixels^16^. Cardioblast nuclei were segmented using a semi-automated approach based on a support vector machine^19^ to detect nuclei, the watershed algorithm^20^ to delineate the detected nuclei, and LiveWire to correct the watershed results. Wound margins were segmented using LiveWire. Epidermal cells were automatically segmented using a seeded watershed algorithm and corrected using LiveWire. Cardioblast nuclei and epidermal cell segmentations were pruned such that all segmented objects were present in every image in the sequence. Tracking was performed by linking objects in adjacent frames by nearest centroid distance^13^.

### Training dataset generation

Wound margin sequences and masks were resized to the network input size (Supplementary Table 1), using bilinear interpolation for image sequences or nearest-neighbour interpolation for masks. Epidermal cell sequences were cropped to the network input size. To train U-Nets and ReSCU-Nets on cardioblast nuclei and wound margin datasets, only every 50^th^ (cardioblast nuclei) or every 5^th^ (wound margins) image in a sequence was used to reduce overfitting due to the similarity between consecutive images. Every image was used in the epidermal cell dataset, as cells changed shape rapidly during the expansion and closure of the wound. LSTM U-Nets required entire videos to train^5^. To train ReSCU-Nets, the mask of each object on the previous frame (for the cardioblast nuclei and epidermal cell datasets) or five frames prior (for the wound margin dataset) was provided as additional input to the network during training. Wound size decreases over time, and thus, we found that training ReSCU-Nets on image/mask pairs five frames apart for the wound margin dataset reduced the dependence of the network on mask information, creating more accurate segmentations during testing.

### Network training

Networks were trained and applied using an NVIDIA RTX 6000 Ada Generation (NVIDIA, Santa Clara, CA) or an NVIDIA TITAN Xp (NVIDIA) graphics processing unit.

Hyperparameters were chosen empirically to produce models of each architecture with the best performance. All networks were implemented in Keras^21^, using the TensorFlow^22^ backend. U-Nets and ReSCU-Nets were trained using the Adamax optimizer^23^ and pixelwise binary cross-entropy loss weighted by the distance transform of the ground truth mask^2^. A random validation split of 10% was used, and an early stopping callback was added to stop training early if the validation loss did not improve, saving the network state with the lowest validation loss upon termination. LSTM U-Nets were trained using the Adam optimizer^5,23^ and weighted pixelwise binary cross-entropy loss^2,5^. A holdout set of one randomly chosen video was used for validation. Training was monitored with TensorBoard^22^, and terminated when the validation loss stopped improving.

### Network training, inference and postprocessing

For datasets with multiple objects in each frame, the networks were prompted with the previous mask of each object separately and trained to segment the corresponding object on the next frame, enabling simultaneous object segmentation and tracking. U-Nets were applied to each image of each test sequence. Following inference, small holes were closed using morphological hole filling. For the epidermal cell dataset, a binary opening was used to separate objects that were connected by small bridges.

LSTM U-Nets were applied to each test sequence, using the entire sequence as the input. Following inference, previously published postprocessing steps were applied to the networks output^5^. Briefly, small holes were filled, and object outlines were enhanced by multiplying the outlines predicted by the network by a mask in which pixels further than 10 pixels from the background were set to 0.

SAM and ReSCU-Nets were applied to each object in each image of the test sequences. The KerasCV implementation of the SA-1B SAM *ViT Huge* was used^8,21^. As prompts, SAM used the bounding box and centroid of the mask for the object in the previous image in the sequence, while ReSCU-Nets used the mask of the object in the previous image. For the first image of a sequence, we used the ground truth of each object to generate prompts for the networks. During inference, the output mask of each object was postprocessed using information already available to the network. For both SAM and ReSCU-Nets, small holes were filled, and false positives were removed. If the highest IOU was below 75%, the ground truth mask of that object was used to prompt the network in the next image. Otherwise, the output mask was used to prompt the network. Following inference, the network masks were postprocessed by a binary hole filling operation applied to each object.

## Code and data availability

We implemented ReSCU-Nets in PyJAMAS^13^, an open-source image analysis tool that we develop. PyJAMAS provides a graphical user interface for training and applying ReSCU-Nets. The source code of PyJAMAS is available at bitbucket.org/rfg_lab/pyjamas/. Additionally, a standalone implementation of ReSCU-Nets is available at <u>bitbucket.org/raymond_hawkins_utor/rescu-net</u>. The datasets used are available in the BioImage Archive^10,24^ at https://www.ebi.ac.uk/biostudies/BioImages/studies/S-BIAD1410.

