## Supplementary Materials for "ReSCU-Nets: recurrent U-Nets for segmentation of multidimensional microscopy data"

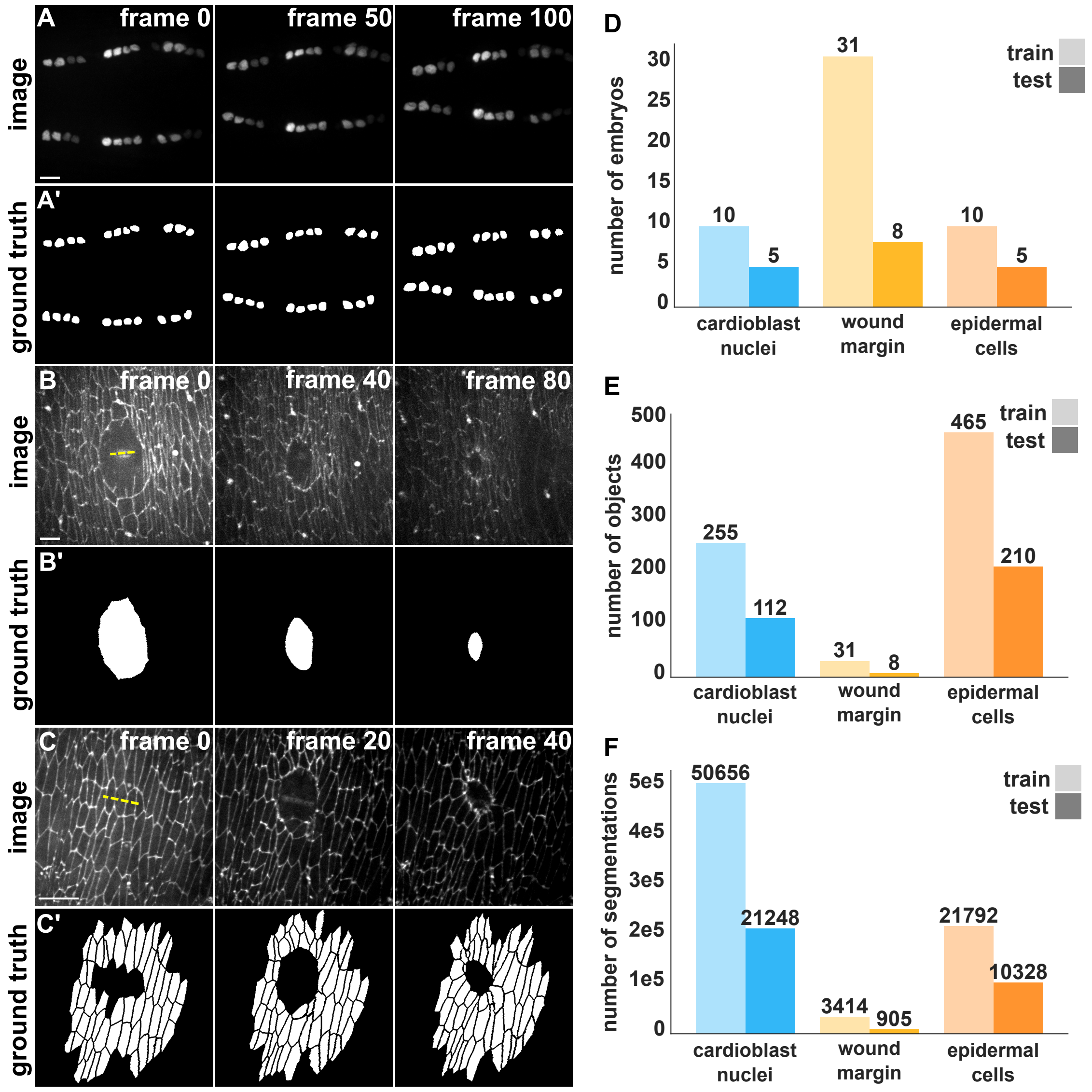


**Supplementary Figure 1. Three timelapse datasets for ReSCU-Net training and testing. (A-C)** Images from time-lapse sequences (A-C) and ground truth segmentations (A'-C') depicting the nuclei of migrating cardioblasts (A), a closing wound (B), and epidermal cells around a wound (C). Yellow dashed lines indicate the region targeted for wounding. Image index in the corresponding sequence is indicated. Anterior, left. Bars, 10 μm. **(D-F)** Number of embryos (D), unique segmented objects (E), and unique object instances (F) for train (light) and test (dark) splits in the cardioblast nuclei (cyan), wound margin (yellow) and epidermal cell (orange) datasets.

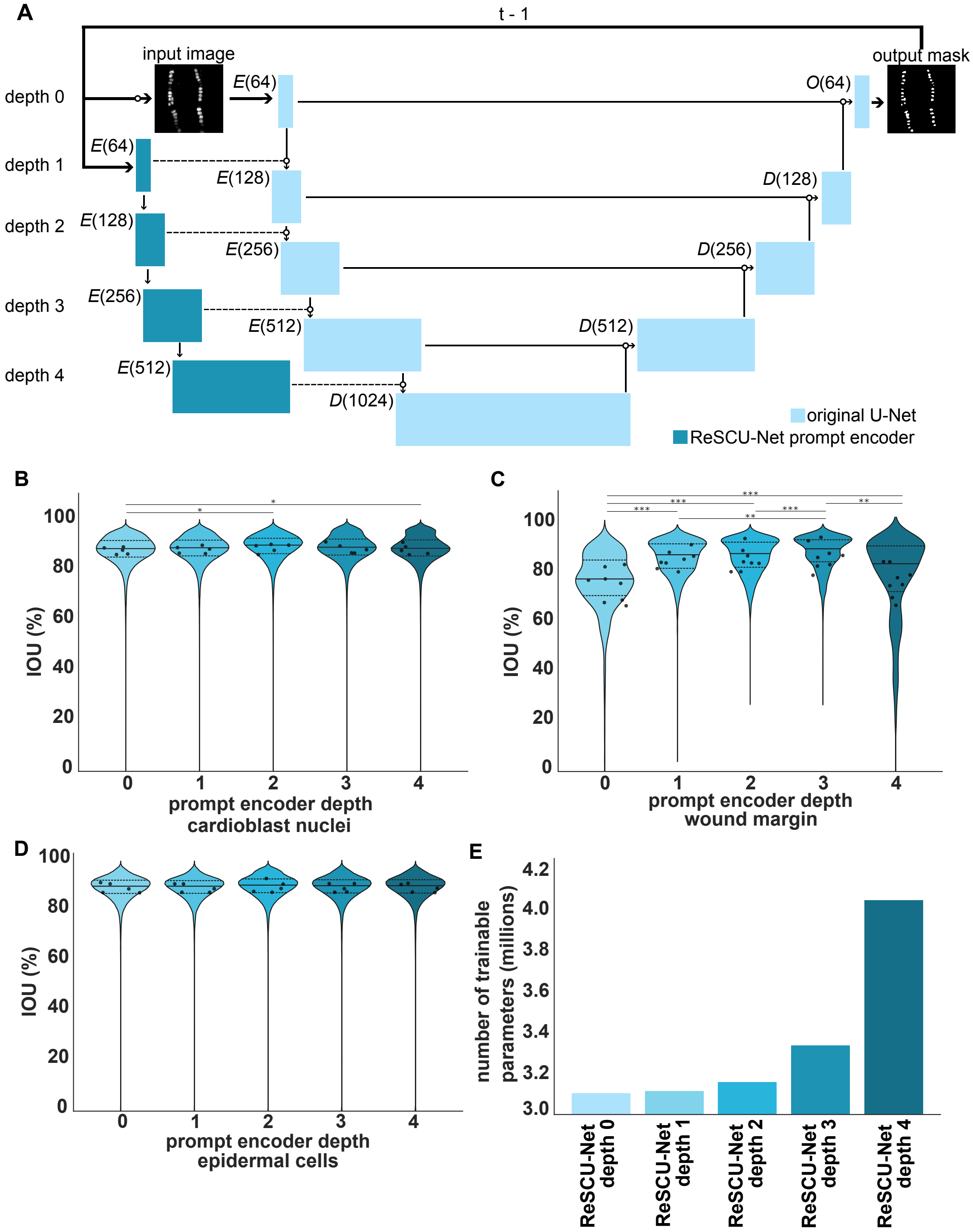

A
t - 1
input image
output mask
depth 0
depth 1
depth 2
depth 3
depth 4
E(64)
O(64)
E(128)
D(128)
E(256)
D(256)
E(512)
D(512)
D(1024)
original U-Net
ReSCU-Net prompt encoder
B
IOU (%)
prompt encoder depth
cardioblast nuclei
C
prompt encoder depth
wound margin
D
prompt encoder depth
epidermal cells
E
number of trainable parameters (millions)
ReSCU-Net depth 0
ReSCU-Net depth 1
ReSCU-Net depth 2
ReSCU-Net depth 3
ReSCU-Net depth 4

**Supplementary Figure 2. ReSCU-Nets with a single mask encoder block provide optimal performance. (A)** ReSCU-Net architecture showing the original U-Net backbone (light blue) and potential ReSCU-Net prompt encoders (dark blue). Dashed lines indicate prompt and image concatenation for each prompt encoder depth. **(B-D)** Test IOUs of ReSCU-Nets with increasing prompt encoder depth for the cardioblast nuclei (B), wound margin (C) and epidermal cell (D) datasets. Violins indicate distributions, solid lines show the median, dashed lines indicate the quartiles. Dots show movie means. * $P < 0.05$, ** $P < 0.01$, *** $P < 0.001$. **(E)** Number of trainable parameters for U-Nets and ReSCU-Nets with increasing recursion depth.

**Supplementary Video 1.** Nuclei of migrating cardioblasts expressing $Mid^{E19}$:GFP, corresponding ground truth segmentation, and segmentations produced by U-Net, LSTM U-Net, SAM, and ReSCU-Net. Cyan and magenta represent pixels erroneously labelled as foreground and background, respectively. Bar, 10 µm. Frames were captured at 15 s intervals.

**Supplementary Video 2.** Closure of a wound in an embryo expressing E-cadherin:tdTomato, corresponding ground truth segmentation of the wound margin, and segmentations produced by U-Net, LSTM U-Net, SAM, and ReSCU-Net. Cyan and magenta represent pixels erroneously labelled as foreground and background, respectively. Bar, 10 µm. Frames were captured at 30 s intervals.

**Supplementary Video 3.** Closure of a wound in an embryo expressing E-cadherin:GFP, corresponding ground truth segmentation of epidermal cells, and segmentations produced by U-Net, LSTM U-Net, SAM, and ReSCU-Net. Cyan and magenta represent pixels erroneously labelled as foreground and background, respectively. Bar, 10 µm. Frames were captured at 30 s intervals.

**Supplementary Table 1.** Hyperparameters for trained networks.

| dataset | heart nuclei | | | wound area | | | cell outlines | | |
|---|---|---|---|---|---|---|---|---|---|
| network | U-Net | LSTM U-Net | ReSCU-Net | U-Net | LSTM U-Net | ReSCU-Net | U-Net | LSTM U-Net | ReSCU-Net |
| image width (pixels) | 512 | 512 | 512 | 192 | 192 | 192 | 256 | 256 | 256 |
| learning rate | $1e^{-3}$ | $1e^{-3}$ | $1e^{-3}$ | $1e^{-3}$ | $1e^{-4}$ | $1e^{-3}$ | $1e^{-3}$ | $1e^{-4}$ | $1e^{-3}$ |
| batch size | 16 | 4 | 16 | 32 | 4 | 32 | 32 | 4 | 32 |
| optimizer | Adamax | Adam | Adamax | Adamax | Adam | Adamax | Adamax | Adam | Adamax |
| concatenation depth | - | - | 1 | - | - | 1 | - | - | 1 |
| unroll length | - | 5 | - | - | 5 | - | - | 5 | - |
